# Regulatory diversity contributes to a divergent transcriptional response to dietary changes in mammals

**DOI:** 10.1101/2020.11.01.364232

**Authors:** Kevin R. Costello, Heon Shin, Candi Trac, Oleg Varlamov, Dustin E. Schones

## Abstract

**BACKGROUND:** Regulatory innovation is central to the evolution of species. Different nutritional sources are one environmental pressure that can lead to selection for novel regulatory elements. Dietary composition changes, including exposure to “western” diets with excess fat and sugar content, can lead to transcriptional regulatory changes in the liver. In order to investigate how transcriptional regulatory changes in response to a high fat diet diverge across species, we profiled chromatin accessibility, histone modifications and the transcriptome in livers of rhesus macaques and mice fed high fat and normal diets.

**RESULTS:** While the majority of elements exhibiting changes in chromatin accessibility in response to a high fat diet are enriched for similar transcription factors across species, the loci that change are mostly species specific. These unique responsive regulatory elements are largely derived from transposable elements and are enriched for liver-specific transcription factors, such as HNF4α. Furthermore, the majority of genes that respond to a high fat diet in rhesus macaques do not have a shared response in mice and are proximal to regulatory elements that display changes in chromatin accessibility only in rhesus macaques.

**CONCLUSIONS:** Our study demonstrates that most of the liver regulatory elements that exhibit changes in chromatin accessibility in response to a high fat diet do so in a species-specific manner. These findings illustrate how a similar environmental stimulus can drive a divergent chromatin and transcriptional responses in evolutionary distinct mammalian species.

## Background

While gene coding regions are highly conserved across mammalian genomes, intergenic regulatory elements tend to be poorly conserved and are a source of phenotypic diversity between mammalian species [1–4]. Both conserved and recently evolved enhancer sequences have been shown to have important phenotypic consequences [4–6]. Highly conserved enhancer sequences can regulate essential pathways such as embryonic development [7]. Novel regulatory elements can arise through spontaneous mutations in DNA sequences, exaptation of ancestral DNA sequences, or though co-option of transposable elements (TE) to drive the establishment of new transcription factor (TF) binding sites (TFBSs) across the genome [8–12]. Novel regulatory elements have been found to play a critical role in modulating the lineage-specific expression of metabolic genes across mammalian species [6,10,12–15]. These novel regulatory elements can allow for species to adapt to and survive new environmental conditions, including dietary changes.

Together with white adipose tissue and skeletal muscle, the liver plays a critical role in regulating glucose and lipid metabolism and systemic insulin sensitivity [15–18]. Although the gene networks involved in hepatic regulation of glucose and lipid metabolism are highly conserved across multiple mammalian species, transcript abundance can be significantly different between species, potentially in response to major dietary shifts, including a transition from vegetarian diets in rodents to meat-containing diets in primates [19–21]. Overconsumption of high-fat diets is associated with ectopic lipid accumulation in the liver, known as non-alcoholic fatty liver disease, and has become a major public health concern in many nations [16,17,22–26]. While the consumption of a high fat diet represents a major obesity risk factor [27], there are additional genetic factors that contribute to and modulate individual responses to a high fat diet [28–30]. Recently, several independent studies have identified novel genetic determinants that contribute to the development of obesity, including gene-specific mutations and changes in the regulatory elements that can potentially alter the expression of genes involved in regulating energy metabolism [31,32]. We have previously reported that long-term exposure to a high fat diet alters the chromatin accessibility state at the gene regulatory regions, resulting in the recruitment of the TFs in the mouse liver [33,34]. These studies demonstrate that dietary factors can induce long-lasting epigenetic changes in the mouse genome and therefore can modulate the expression of the key TFs involved in the regulation of energy metabolism in the liver.

While the mouse has become a widely accepted model system for studying obesity, there are few human studies investigating the impact that diet-induced obesity can have on the epigenetic and transcriptional landscape as these studies are difficult to conduct because of the confounding effects. So to bridge the gap between rodent and human studies we utilized a nonhuman primate model of diet-induced obesity [35–38]. In the present study, we profile the chromatin accessibility and transcriptional responses of rhesus macaque livers prior to and after high fat feeding [36,37] and compare these responses to those from liver data sets derived from mice fed a standard chow or a high fat diet [33,39]. Our analysis reveals that the rhesus macaque and mouse genomes contain divergent sets of regulatory elements that respond to a high fat diet, contributing to species-specific transcriptional changes in response to a high fat diet.

## Results

### Identification of conserved and divergent changes in chromatin accessibility between mice and rhesus macaques in response to a high fat diet

To determine the dietary regulation of chromatin accessibility and transcription in intact adult rhesus macaques, we performed the ATAC-seq, ChIP-seq, and RNA-seq analyses using rhesus macaque liver biopsies that were collected before and after the animals were fed a high fat diet for a 6 month period [36,37] (Fig. 1a). The dietary fat content changed from 15% to 33% when these monkeys were put on a high-fat diet [37]. All monkeys profiled exhibited a significant increase in body fat after 6 months of high fat feeding (2.93 ± 0.26-fold increase) (Additional file 1: Fig. S1a, Additional file 2) [37]. In order to identify shared and species-specific changes, we also utilized chromatin accessibility data from the livers of mice fed either a standard chow (10% fat) or high fat (30% fat) diet for 8 weeks [29]. As we were interested in profiling changes in chromatin accessibility driven by a high fat diet, we only focused on regulatory elements that have similar changes in chromatin accessibility in response to high fat diet between individual rhesus macaques (Additional file 1: Fig. S1b).

**Fig. 1.**
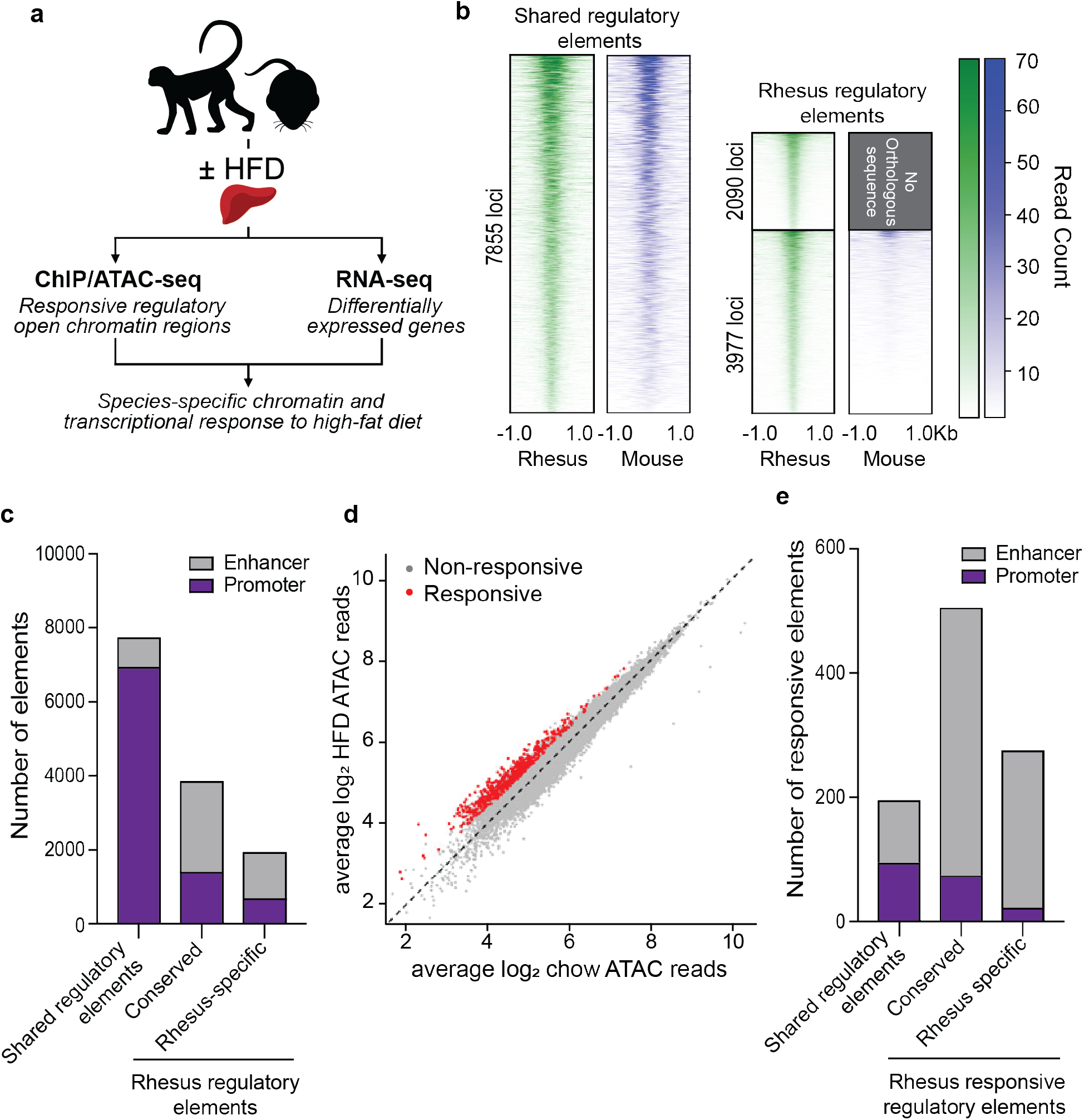
Regulatory elements that are unique to rhesus macaques are the most responsive to a high fat diet. **a** Schematic of experimental design. **b** Heatmaps showing chromatin accessibility for all regulatory elements found in rhesus macaques (green) and the orthologous loci in mice (blue). Regulatory elements with no orthologous sequence in the mouse genome are displayed in grey. Shared regulatory elements were loci found to be accessible in both rhesus macaques and mice. Rhesus regulatory elements were loci that were only accessible in rhesus macaques. **c** Breakdown of regulatory elements into promoters or enhancers based on H3K4me3 or H3K27ac enrichment, respectively. Regulatory elements that are only accessible in rhesus macaques were further broken down into loci that have any sequence conservation to the mouse genome or are rhesus-specific elements with no underlying sequence conservation. **d** Identification of the top 1000 most responsive regulatory elements in response to a high fat diet. **e** The number of promoter and enhancer elements that respond to a high fat diet in rhesus macaques and mice, or those that respond only in rhesus macaques.

In order to determine the extent to which the orthologous loci identified in rhesus macaques were also accessible in mice, we mapped all of the regulatory elements from the rheMac10 to mm10 genome using LiftOver. Of the 13,922 putative regulatory elements we identified in rhesus macaques, 7,855 (56%) were determined to be shared regulatory elements that were accessible in both rhesus macaque and mouse, while the remaining 6,067 (44%) were found to be rhesus regulatory elements, meaning they were only accessible in rhesus macaque (Fig. 1b). We further subdivided the rhesus regulatory elements into those that were conserved with some underlying sequence conservation to the mouse genome (65.6% of loci) or those that were rhesus-specific with no sequence conservation between the two genomes (34.4% of loci). Using H3K4me3 ChIP-seq enrichment to annotate promoters and H3K27ac enrichment to annotate enhancers, we found that promoter elements were more likely to be accessible in both genomes, while enhancer elements were more likely to be unique to the rhesus macaque genome (Fig. 1c), consistent with previous work [40].

To identify the regulatory elements that were the most responsive to dietary changes, we selected the top 1000 regulatory elements that displayed the greatest increase in chromatin accessibility in response to a high fat diet (Fig. 1d; Additional file 1: Fig. S1b). We found a small fraction (22%) of responsive regulatory elements are accessible in both genomes. The majority of these responsive elements that are accessible in both genomes demonstrate similar changes in chromatin accessibility in response to a high fat diet (Additional file 1: Fig. S1c), indicating that these shared regulatory elements have a conserved function between species. However, we found that the majority of regulatory elements (78%) which respond to high fat diet are unique to rhesus macaques (Fig. 1e), demonstrating that mice and rhesus macaques have evolved a divergent response to dietary changes. Together this work demonstrates divergent changes in chromatin accessibility in response to a high fat diet.

### Liver transcription factors drive changes in chromatin accessibility in response to a high fat diet

As previous studies in mice have shown transcription factor recruitment is responsible for driving changes in chromatin accessibility in response to a high fat diet [33], we profiled TF motifs enriched in rhesus macaque responsive regulatory elements. We found that binding motifs for liver TFs, including HNF4α, FOXA1 and CEBPα, were significantly enriched at responsive regulatory elements in rhesus macaques (Fig. 2a, Additional file 1: Fig. S2a). These TFs were of significant interest to us as they were shown to drive changes in chromatin accessibility in mice after high fat diet feeding and bind cooperatively to regulate transcription [33,41]. Using existing ChIP-seq datasets for HNF4α, FOXA1 and CEBPα from rhesus macaque livers [2], we determined that nearly all responsive regulatory elements are cooperatively bound by these TFs (Fig. 2b).

**Fig. 2.**
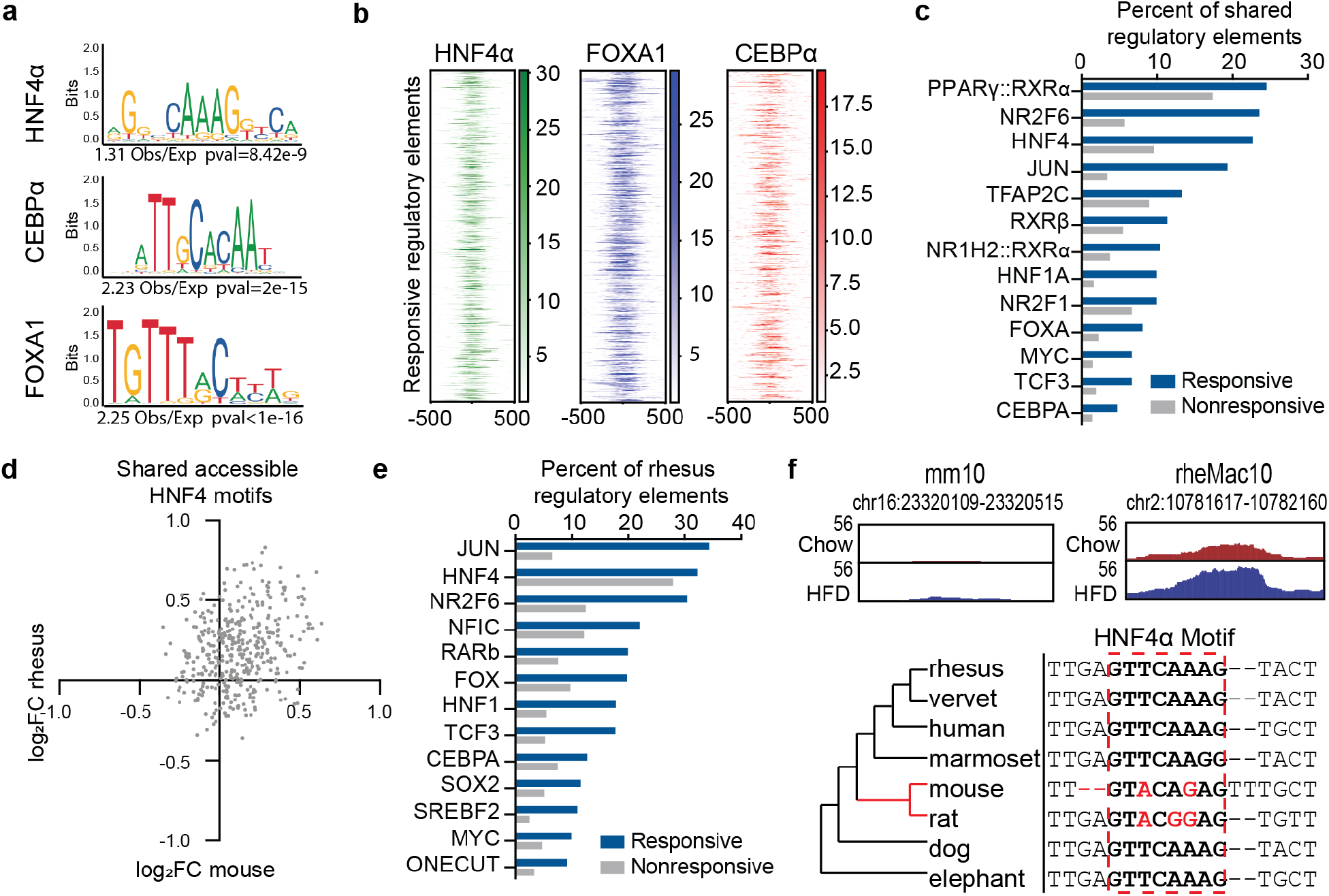
Liver transcription factors drive changes in chromatin accessibility in response to a high fat diet. **a** Enrichment of binding motifs for HNF4α, FOXA1 and CEBPα in responsive regulatory elements. Significance was determined using a chi-squared test to compare enrichment of TFBS at the responsive regulatory elements relative to the non-responsive elements. **b** Profile of TF occupancy by ChIP-seq at responsive regulatory elements. **c** Identification of TF binding motifs on shared responsive regulatory elements that are conserved between both mice and rhesus macaques. **d** Profile of chromatin accessibility changes in response to HFD at orthologous loci containing a conserved HNF4α binding motif. **e** Identification of novel TF binding motifs on responsive REs that are unique to rhesus macaques. **f** Top: an example of a regulatory element that is responsive to a high fat diet in rhesus macaques but not in mice. Bottom: multiple sequence alignment highlighting mutations in the HNF4 binding motif (boxed by the red dotted line) in the mouse genome.

Given that recently evolved regulatory elements potentially have different transcription factor binding sites than conserved regulatory elements [2,8,40], we were interested in profiling whether similar TFs were responsible for driving changes in chromatin accessibility at both shared and rhesus responsive regulatory elements. We profiled the shared regulatory elements to identify all TF binding motifs present in both rhesus macaques and mice and examined which of these TFBSs were enriched in the responsive fraction. HNF4α, FOXA1, and CEBPα binding motifs were among the top motifs that were found to be responsive to high fat diet and conserved between mice and rhesus macaques at the shared regulatory elements (Fig. 2c). Shared regulatory elements that contain HNF4α binding motifs in both mice and rhesus macaques have a similar increase in chromatin accessibility in response to high fat diet for both rhesus macaques and mice (Fig. 2d), indicating a shared function for these elements in both species. This is in contrast to SP1 motifs, where there is little to no change in chromatin accessibility in response to a high fat diet in both species (Additional file 1: Fig. S2b). This demonstrates that these shared responsive regulatory elements are functionally conserved due to retention of TFBSs between mice and rhesus macaques.

We also determined whether responsive regulatory elements that are unique to rhesus macaques have novel TF binding sites that can be responsible for driving rhesus specific changes in chromatin accessibility. Novel HNF4α, FOXA1, and CEBPα binding motifs were found to be enriched at the unique responsive regulatory elements in rhesus macaques, as well as motifs for other TFs such as JUN and NR2F6 (Fig. 2e) that have been shown to be essential for the regulation of metabolism in the liver [42,43]. At loci with partial sequence conservation, these novel TFs binding motifs can be the result of sequence mutations in TF recognition sequences driving the divergence of regulatory element response between species (Fig. 2f). Together this work demonstrates that while the same transcription factors are responsible for changes in chromatin accessibility in both mice and rhesus macaques, the majority of responsive loci are unique to the rhesus macaque genome.

### Transposable elements provide novel responsive regulatory elements through the propagation of liver transcription factor binding sites

Given that transposable elements are known to be co-opted as regulatory elements in mammals [8,11,40], we sought to examine the extent to which TEs contribute to the formation of regulatory elements that respond to a high fat diet in rhesus macaques. We observed significant enrichment of TEs at responsive regulatory elements that are unique to rhesus macaques (Fig. 3a). As previous work has demonstrated that exaptation of older TEs can help drive novel regulatory elements [40], we worked to determine if these TE-derived responsive regulatory elements were the result of co-option of old or young TEs. We profiled the evolutionary age of TEs found at responsive regulatory elements using the sequence divergence of each element from the consensus TE sequence as a proxy for age [11]. TEs were separated into two groups: young TEs that have a sequence divergence less than 2 percent (or a millidiv of 20) which are mostly unique to primates, and old TEs that have a sequence divergence greater than 2 percent (or a millidiv of 20) which are found in both primates and rodents (Fig. 3b). We found that 67% of responsive regulatory elements are the result of the co-option of older TEs (Fig. 3b).

**Fig. 3.**
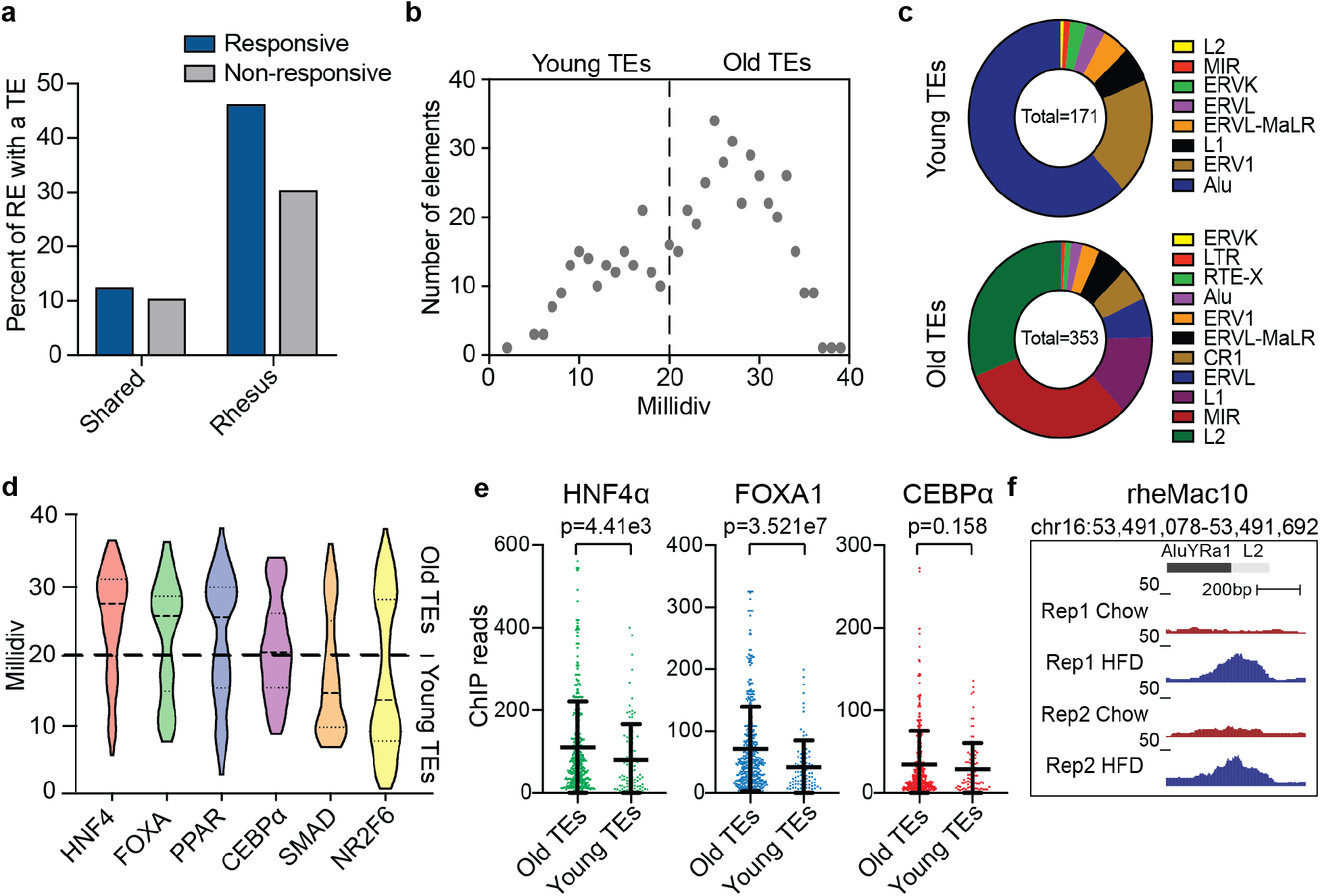
Transposable elements drive the formation of unique responsive regulatory elements through the propagation of transcription factor binding sites. **a** Percent of regulatory elements that contain a transposable element. **b** The number of TEs found on responsive regulatory elements separated by sequence divergence. **c** Breakdown of young (milliDiv < 20) and old (milliDiv > 20) TE families found at responsive regulatory elements. **d** Presence of responsive TF binding motifs on all TE-containing regulatory elements as determined by motif scanning using FIMO. TEs were separated by sequence divergence (miliDiv) from consensus as an estimate of age. **e** ChIP-seq signal intensity at responsive regulatory elements containing either old or young TEs. Significance determined by t-test. **f** An example of a responsive regulatory elements that is derived from a TE and is unique to rhesus macaques

We observed a strong TE family bias, where responsive younger TEs are nearly all Alu and ERV1 elements, while the older TEs are mainly L2 or MIR elements (Fig. 3c). Some TE families, such as L2s and ERV1s, are present in both the young and old fraction since individual elements may have a sequence divergence that is higher or lower than the 20 millidiv divergence. These young and old TEs have distinct TF binding motif profiles. Older TEs are enriched for binding motifs of liver TFs such as HNF4α and FOXA1, while younger TEs have enrichment for other TF binding motifs such as NR2F6, which is an inflammatory TF (Fig. 3d). These results support our previous work demonstrating how old TEs display enrichment of liver TFs such as HNF4α [8] while younger TEs have enrichment of inflammatory TF binding sites [12]. Indeed, by profiling ChIP-seq signals for HNF4α, FOXA1 and CEBPα at responsive regulatory elements containing old and young TEs, we found that HNF4α and FOXA1 appear to have a strong bias toward the older TEs, while CEBPα seems to be bound to both young and old TEs, confirming the results found by motif scanning (Fig. 3e). This co-option of TEs can drive the expansion of novel TF binding sites across the genome that can respond to dietary changes (Fig. 3f), highlighting how co-option of transposable elements can aid in the propagation of novel regulatory elements that can respond to environmental changes.

### Novel responsive regulatory elements drive species-specific transcriptional responses to high fat diet

Given that we observed both shared and divergent changes in chromatin accessibility between mice and rhesus macaques, we worked to identify the shared and divergent changes in gene expression as well. We used RNA-seq to profile the transcriptomes from control and high fat-fed rhesus macaques and complemented this with RNA-seq data from mice fed standard chow and high fat diets [39]. Of the transcripts that were differentially expressed in rhesus macaques in response to a high fat diet, approximately one-fifth had a shared transcriptional response in mice (Fig. 4a). As expected, the transcripts that had a shared response to a high fat diet are mainly involved in the regulation of metabolic processes (Fig. 4b,c). We next worked to determine if these shared responsive transcripts could be regulated by the shared responsive regulatory elements. We found an enrichment of responsive shared regulatory elements proximal (<500kb) to these shared responsive transcripts (Fig. 4d). An example of a gene with a shared response to high fat diet is *MFSD2A* (Fig. 4e,f), which is a known regulator of metabolism in the liver [44,45]. This gene is proximal to a shared responsive regulatory element (Fig. 4f), demonstrating functional conservation between rhesus macaques and mice in response to a high fat diet.

**Fig. 4.**
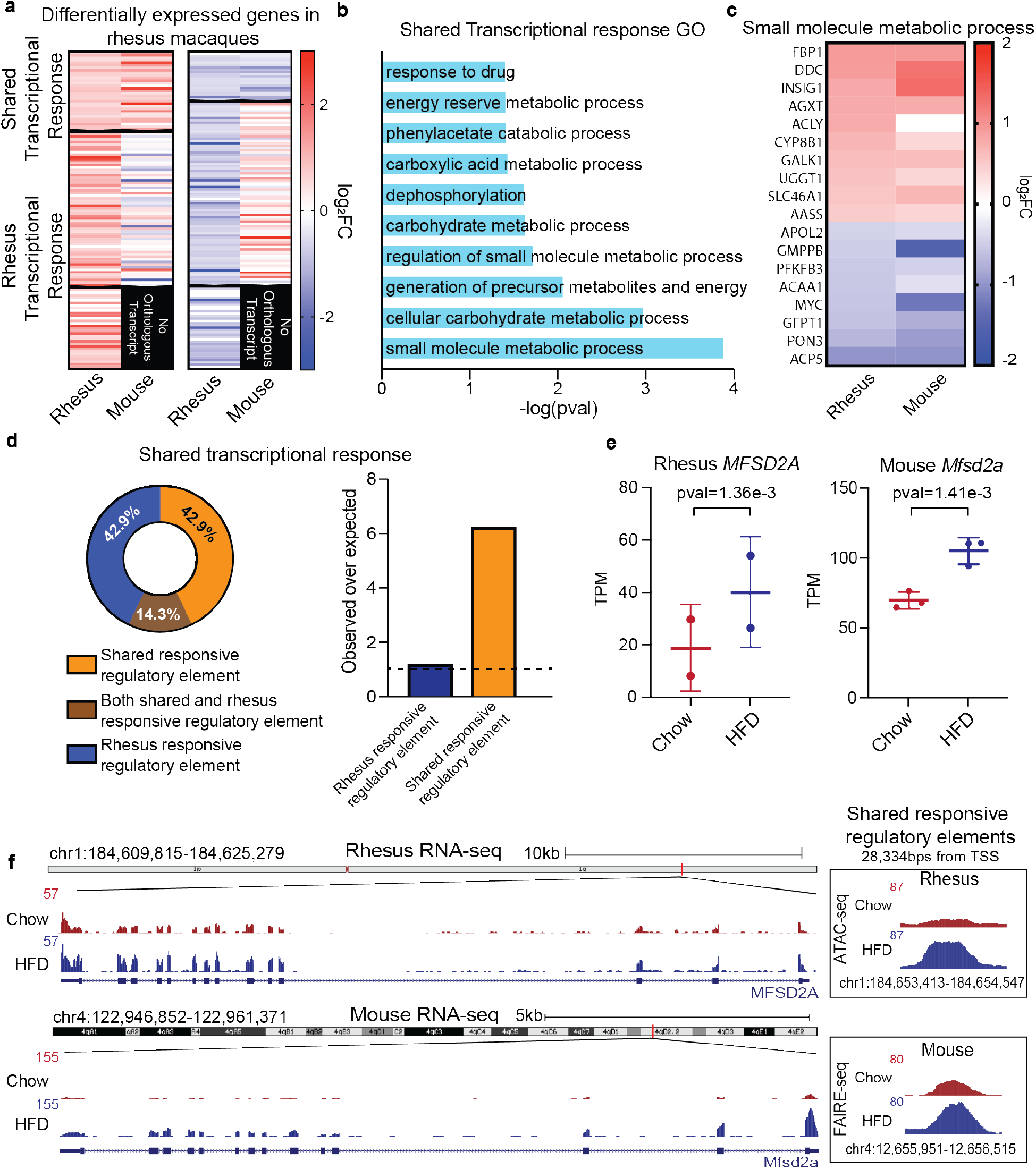
Shared responsive regulatory elements drive a shared transcriptional response to a high fat diet in rhesus macaques. **a** Heatmap of transcripts that have a significant *(FDR* < 0.1) increase (left) or decrease (right) in expression in rhesus macaques in response to high fat diet. **b** Gene ontology of the top 10 gene pathways impacted by high fat diet in both rhesus macaques and mice **c** fold change of genes involved in small molecule metabolism that have a shared response to high fat diet. **d** Enrichment (left) and distribution (right) of shared and rhesus specific regulatory elements within 500kb of TSS shared responsive genes. **e** Fold change in response to high fat diet for both rhesus and mice of the gene *MFSD2A* that is differentially expressed in both mice and rhesus macaques. **f** Genome screenshot of RNA-seq (left) and chromatin accessibility (right) of *MFSD2A* and the shared responsive regulatory element proximal to its TSSs in both rhesus macaques (top) and mice (bottom).

As the majority of transcripts that respond to a high fat diet are uniquely responsive in rhesus macaques, we sought to determine what contribution shared and rhesus specific regulatory elements made to the transcriptional response observed. We found that rhesus responsive transcripts were also involved in regulation of metabolic processes (Fig. 5a,b). We next asked if these rhesus responsive transcripts could be regulated by the rhesus responsive regulatory elements. We found enrichment of rhesus responsive regulatory elements proximal (<500kb) to these rhesus responsive transcripts (Fig. 5c). Interestingly, we found the metabolic gene, glucose-6-phosphatase catalytic subunit *(G6PC)* [46,47] as an example of a transcript with a unique response to a high fat diet between rhesus macaques and mice (Fig. 5d,e). This transcript is proximal to an enhancer with a change in chromatin accessibility that is unique to rhesus macaque genome. This regulatory element has mutations in sequence that allow for primate-specific recruitment of FOXA1 (Fig. 5f). This work demonstrates how divergence regulatory landscape through the deposition and disruption of TF binding sites can impact the transcriptional response of rhesus macaques to a high fat diet.

**Fig. 5.**
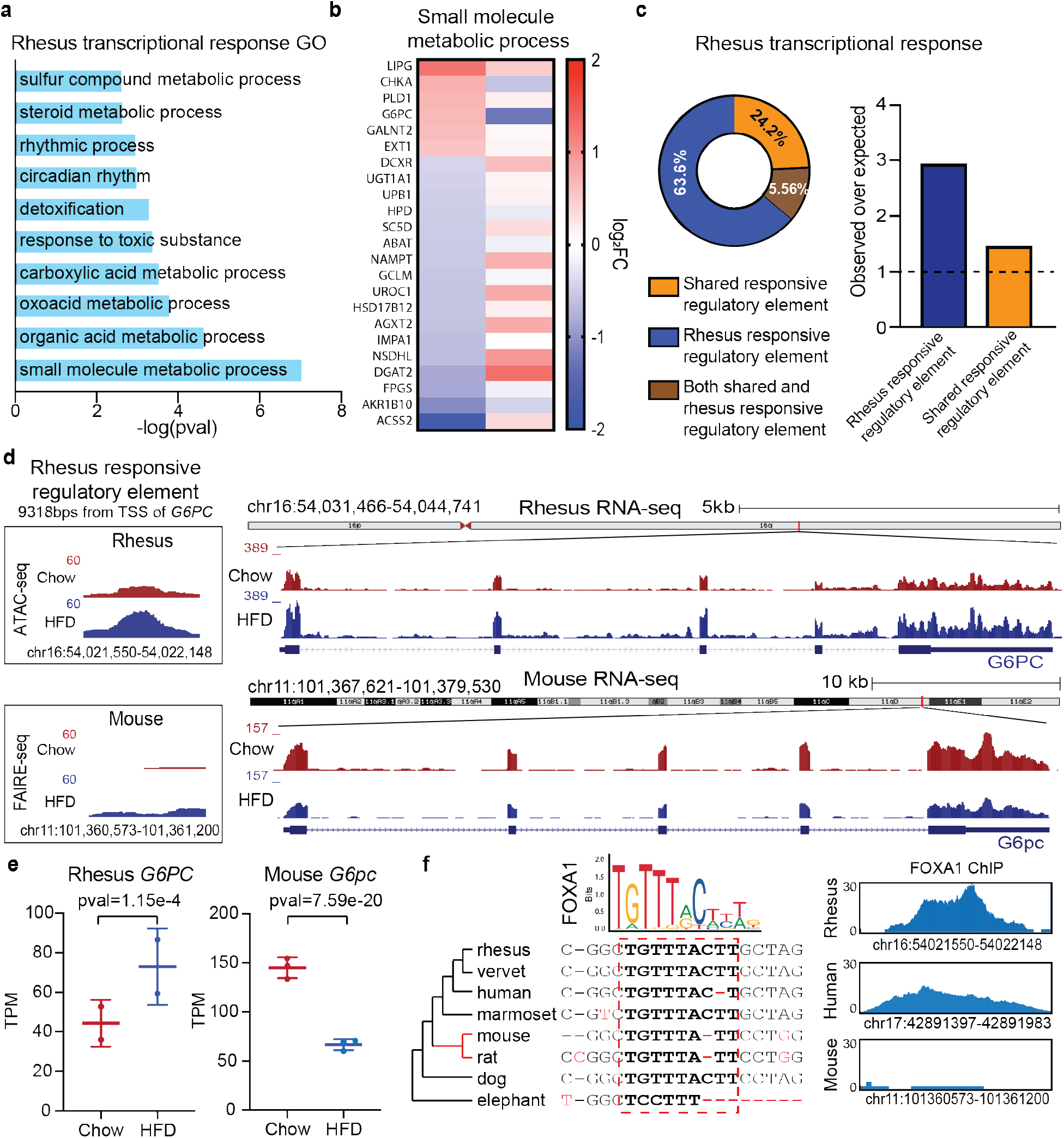
Divergent transcriptional response to a high fat diet between rhesus macaques and mice impact different gene pathways. **a** Gene ontology of the top 10 gene pathways impacted by high fat diet in both rhesus macaques and mice. **b** Fold change of genes involved in small molecule metabolism that have a response to high fat diet that is unique to rhesus macaques. **c** Enrichment (left) and distribution (right) of shared and rhesus specific regulatory elements within 500kb of the TSS of genes that have response to high fat diet that is unique to rhesus macaques. **d** Genome screenshot of RNA-seq (right) and chromatin accessibility (right) of the gene *G6PC* and the rhesus responsive regulatory element proximal to its TSSs in both rhesus macaques (top) and mice (bottom). **e** Fold change in response to high fat diet for both rhesus and mice of the gene G6PC. **f** Multiple sequence alignment of the rhesus responsive regulatory element domestrating FOXA1 binding motif that is not found in mice (left). FOXA1 ChIP signal at orthologous regions in rhesus, humans, and mice (right).

## Discussion

Mammals have evolved to survive fluctuations in the availability and nutritional content of food sources. During periods of overnutrition, metabolic adaptations occur in the liver that promote the storage of metabolic precursors, such as triglycerides [48]. While such a response has been evolutionarily advantageous for mammals, as food supply was not always in abundance, the spread of the high-fat, high-calorie western style diet in humans has contributed to the increasing incidence of both obesity and non-alcoholic fatty liver disease throughout the world [16,17,22–26]. Although the mouse has become a widely accepted model system for studying obesity, major dietary shifts have contributed to altered regulation of metabolic genes between mice and primates [19–21]. Due to this divergence in diet, we make use of a rhesus macaque obesity model that we have developed in order to profile the liver chromatin and transcriptional response to a high fat diet, and compare these changes to those that occur in mice in order to highlight the divergent evolutionary response between these two species to a metabolic stress.

We have previously demonstrated that high fat diet feeding in mice leads to changes in the chromatin accessibility of enhancer elements [12,33,34]. Through our comparative analysis of chromatin accessibility changes between mice and rhesus macaques, we found that the majority of elements that respond to a high fat diet are unique to a particular species. Interestingly, we observed that the same liver TFBSs are found at responsive regulatory elements in both mice and rhesus, including HNF4α, FOXA1 and CEBPα [33]. However, the majority of these binding sites are not conserved between rhesus macaques and mice. These novel transcription factor binding sites can be formed through sequence mutations or by co-option of transposable elements to propagate transcription factor binding sites. Finally, we observed that these unique responsive regulatory elements can potentially drive a unique transcriptional response to high fat diet (Fig. 6), demonstrating how divergent changes in chromatin accessibility can impact neighboring gene expression.

**Fig. 6.**
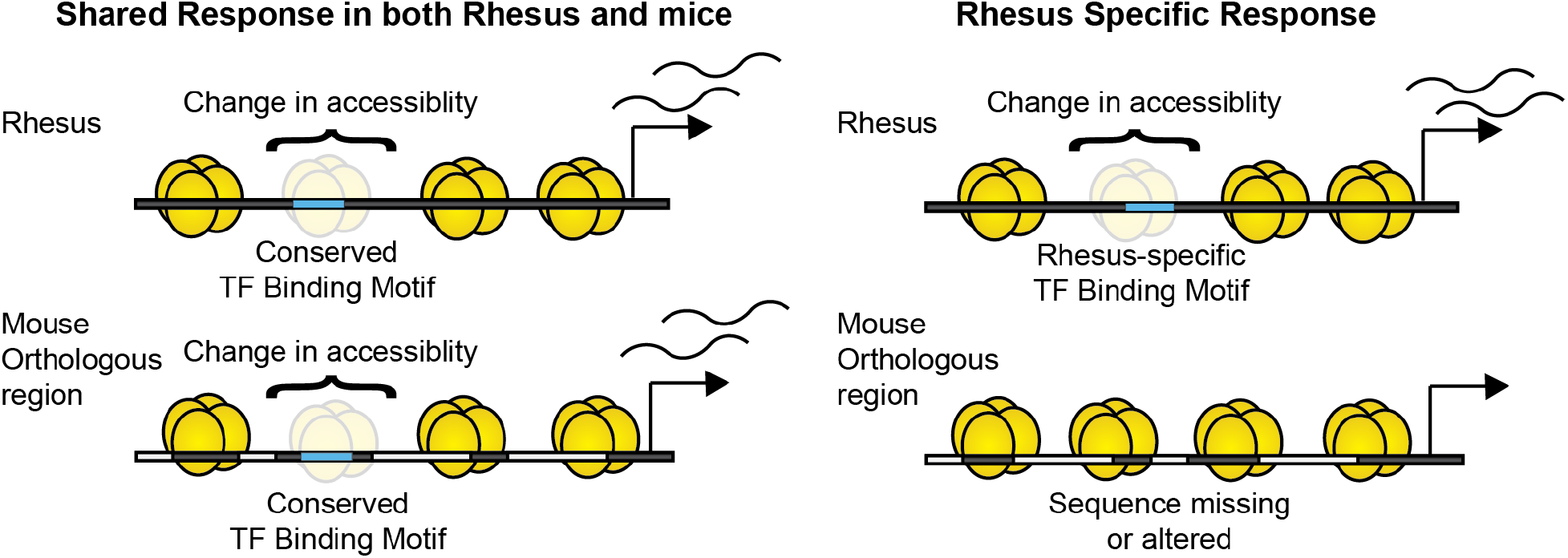
Model of species-specific transcriptional regulation though novel regulatory elements in response to a high fat diet. Shared responsive regulatory elements have conservation transcription factor binding motifs and can drive a shared transcription response to high fat diet. Unique responsive regulatory elements can have novel transcription factor binding sites and drive a species-specific response to high fat diet. can potentially drive a unique transcriptional response to high fat diet

Our study shows that a subset of genes controlling carbohydrate and lipid metabolism in response to a high fat diet exhibit strikingly different transcriptional regulation in mice and nonhuman primates. For example, the *G6PC* gene was significantly upregulated by a high fat diet in the macaque liver but not in the mouse liver. Given the reported involvement of this gene in controlling gluconeogenesis, hepatic glucose output, and its role in the pathogenesis of nonalcoholic fatty liver disease [49,50], we speculate that the transcriptional regulation of hepatic glucose output is more beneficial to larger mammals, such as nonhuman primates and humans, while smaller mammals may control hepatic glucose output by more posttranscriptional mechanisms. Future studies can help understand whether the interspecies differences in the transcriptional regulation of *G6PC* reported in our study contribute to the species-specific pathology of non-alcoholic fatty liver disease in mice and humans. We anticipate that nonhuman primates (and possibly humans) are more susceptible to non-alcoholic fatty liver disease than mice in part because the gene of Fatty Acid Binding Protein 1 *(FABP1)* [51] is positively regulated by a high fat diet in macaques but not in mice (Additional file 1: Fig. S3a,b). Consistent with this hypothesis, a previous study has shown that the overexpression of *FABP1* in the mouse liver promotes non-alcoholic fatty liver disease [52].

It is well understood that elements that are highly conserved between species play essential biological roles. This work, however, suggests that the majority of regulatory elements that respond to a high fat diet are in fact novel enhancer elements. A future direction for this work would be to define this response to high fat diet in other species, to determine exactly how dynamic these responsive regulatory elements are between species, and at what rate novel elements are arising. It would also be interesting to elucidate if other environmental stimuli, such as stress and exercise, can be impacted by species-specific changes in chromatin accessibility.

## Conclusion

Our results indicate a divergent regulatory and transcriptional response to high fat diet between rhesus macaques and mice. We found the majority of elements that respond to a high fat diet are unique to rhesus macaques. Interestingly, while similar TFs drive response in both rhesus and mice, the loci where these TFs are being recruited are mostly species specific. Additionally, the majority of genes with significant changes in transcription are proximal to responsive open chromatin sites. Altogether, this work demonstrates how an environmental stimulus, such as exposure to a high fat diet, results in a divergent chromatin and transcriptional response between two mammals.

## Methods

### Animals

Rhesus macaque liver samples were collected from animals described in previous studies [36,37]. Briefly, 12-13 month old animals were given 2 months of Standard chow diet, consisting of the two daily meals of Purina Lab Diet fiber-balanced monkey chow (15% calories from fat, 27% from protein, and 59% from carbohydrates; no. 5000; Purina Mills, St. Louis, MO), supplemented with fruits and vegetables. Animals were then given a high fat diet (33% calories from fat, 17% from protein, 51% from carbohydrates; 5A1F, Purina Mills, St. Louis, MO) ad libitum for 6 months. Liver biopsies were collected longitudinally before and after animals were exposed to a high fat diet. Food was withheld for approximately 12 hrs prior to the procedure. Water was not withheld. Animals were sedated with 8-20 mg/kg ketamine administered intramuscularly. An intravenous catheter was placed, and the animal received 0.025-0.2 mg/kg hydromorphone HCl intravenously. Following sedation a local anesthetic block consisting of 0.4 ml bupivicaine (0.5%) was administered intradermally at the incision site. Animals were intubated with an appropriate size endotracheal tube and a surgical-depth of anesthesia was maintained with 0.5-2% isoflurane. Isoflurane will be combined with 100% oxygen administered at a rate of 1-1.5 L/min. Animals were placed on mechanical ventilation for all laparoscopic procedures while the abdomen is insufflated with CO2. The ventral abdomen was shaved from the xiphoid process to the pubis and laterally to the flank folds. A sterile prep using ChlorPrep (a chlorhexidine/alcohol instant solution) was applied to the entire surgical site and allowed to dry. All surgical procedures were conducted by trained Surgical Services Unit personnel under the supervision of surgical veterinarians in dedicated surgical facilities using aseptic techniques and comprehensive physiological monitoring. Animals received intravenous fluids (generally Lactated Ringers Solution) at 5-20 ml/kg/hour. The animals were positioned in dorsal or slight right lateral recumbency followed by sterile preparation and draping of the abdomen. A Verres needle was inserted via 5 mm skin incision on the abdomen to the left of the umbilicus followed by insufflation to 8-15 mm Hg pressure with CO2 gas. The Verres was removed and the 5 mm trocar/sheath and 5 mm telescope was inserted by puncture at the same site. A left paracostal 5 mm accessory port was placed, through which a cutting biopsy grasper is inserted. Two pinch biopsies were removed from the caudal margin of the left/right liver lobe. Hemostasis was verified by direct observation and assisted by direct pressure with a cotton swab if needed. The laparoscopic instruments were removed and the incisions were closed with interrupted 4-0 Monocryl in the rectus fascia and skin. Animals underwent the postsurgical recovery and the liver biopsies were repeated several months later following a high fat diet period, as described [36].

### ATAC-seq

Chow and high fat diet fed monkeys biopsied livers were used for ATAC-seq (Assay for Transposase-Accessible Chromatin using sequencing) as described previously [53]. Isolated DNA was sequenced on an Illumina HiSeq 2500 System. We assessed standard QC measures on FASTQ file http://www.bioinformatics.babraham.ac.uk/projects/fastqc and adapters were trimmed using Trimgalore http://www.bioinformatics.babraham.ac.uk/projects/trim_galore/. Trimmed reads were aligned to the RheMac10 genome using bowtie1 retaining only reads that could be mapped to unambiguously to a single location in the genome. Aligned reads were sorted using samtools [54] and filtered for duplicate reads using the MarkDuplicates function of Picard tools http://broadinstitute.github.io/picard/.

### ChIP-seq

Chromatin immunoprecipitation was performed with H3K27ac antibody (ab4729, Abcam), H3K4me3 antibody (ab8580, Abcam) and IgG control (ab37415, Abcam) as previously described [55]. Isolated DNA was sequenced on an Illumina HiSeq 2500 System. We assessed standard QC measures on FASTQ file using FASTQC http://www.bioinformatics.babraham.ac.uk/projects/fastqc and adapters were trimmed using Trimgalore http://www.bioinformatics.babraham.ac.uk/projects/trim_galore/. Trimmed reads were aligned to the RheMac10 genome using bowtie1 retaining only reads that could be mapped to unambiguously to a single loci. Aligned reads were sorted [54] and filtered for duplicate reads using the MarkDuplicates function of Picardtools http://broadinstitute.github.io/picard/. Regions of histone enrichment were called using MACS2 callpeaks [56] with a *q*-val threshold of 0.001. the -broad option was added for profiling H3K27ac ChIP data. Only peaks found in both monkeys were retained to remove biological noise.

### RNA-seq

RNAs were extracted from biopsied monkey livers using Trizol (Invitrogen) and RNA was polyA enriched. Eluted RNAs were sequenced by Illumina HiSeq 2500 System. We assessed standard QC measures on FASTQ file http://www.bioinformatics.babraham.ac.uk/projects/fastqc and adapters were trimmed using Trimgalore http://www.bioinformatics.babraham.ac.uk/projects/trim_galore/. Trimmed reads were aligned to the RheMac10 genome using STAR [57]. Coverage of both Rhesus transcripts were determined using Bedtools coverage [58] and variability caused by interindividual differences were corrected using limma [59]. Differentially expressed genes were identified using Deseq2 as genes with an FDR less than or equal to 0.1 [60]. Gene ontology enrichment analysis was performed using PANTHER [61–63].

### Profiling of ATAC-seq data

Loci of accessible chromatin were identified using MACS2 callpeaks [56] with a *q*-val threshold of 0.001. Only loci which were found to be accessible in both monkeys were retained to remove biological noise. Total read numbers mapping to open chromatin loci was calculated using Bedtools coverage [58] and normalized by reads mapping to peaks. As we were interested in profiling changes in response to a high fat diet, elements that had a log2FC difference greater than 0.50 between both monkeys were filtered out, to remove all elements that may be responding as a result of biological noise. Additionally, all open chromatin loci found to not be at a region of H3K27ac or H3K4me3 enrichment were disregarded from the analysis. The top 1000 elements with the greatest increase in chromatin accessibility were selected as our responsive regulatory elements. All regulatory elements were mapped to from the rhesus macaque genome to the mouse genome using liftOver [64] with the -minMatch=0.1 and -multiple allowing loci to map multiple locations and prevent artificall overrepresentation of Tes in the rhesus unique fraction.

### Motif scanning

Motif scanning was performed using FIMO [65] to scan for vertebrate motifs from the JASPAR database [66]. All motifs with a *p*-val > 0.001 were retained for further analysis. Conserved and rhesus-specific motif profiling was performed by comparing the presence of all TF binding motifs at accessible loci in the rhesus macaque genome. If the motif was found to be in both rhesus and mice, it was determined to be conserved, while if it was only found in rhesus it was determined to be a rhesus specific motif. Additionally, results for motifs with similar profiles (i.e. HNF4α and HNF4γ) were merged to prevent redundancy and improve clarity for Figure 2.

## Abbreviations

TF: Transcription factor,
TFBS: Transcription factor binding site,
ATAC: Assay for Transposase-Accessible Chromatin sequencing,
ChIP-seq: Chromatin immunoprecipitation sequencing,
RNA: Ribonucleic acid sequencing,
HFD: High fat diet,
HNF4α: Hepatocyte nuclear factor 4 alpha,
C/EBPα: CCAAT/enhancer binding protein alpha,
FOXA1: Forkhead box A1,
STAT: Signal transducers and activators of transcription.

## Supplementary information

## Acknowledgements

We thank Dr. Amy Leung and members of the Schones lab for helpful comments and suggestions.

## Funding

Funding was provided by the National Institutes of Health, grants R01DK112041 and R01CA220693 (D.E.S.) and R21 AG047543 (O.V.). Research reported in this publication included work performed by Integrative Genomics Cores of the City of Hope supported by National Institutes of Health under NCI Award P30CA33572. The authors declare that they have no conflicts of interest with the contents of this article.

## Availability of data and materials

All datasets generated in this study have been submitted to the NCBI Gene Expression Omnibus (GEO; https://www.ncbi.mln.nih.gov/geo/) under accession no. GSE160247.

Mouse chromatin accessibility data was from GEO:GSE55581. HFD RNA-seq data was from GEO:GSE77625. Rhesus macaque ChIP-seq data for HNF4, CEBPa, and FOXA1 are from arrayExpress E-MTAB-1509. Human FOXA1 and HNF4a ChIP-seq data is from Encode project, at GEO:GSE91618 and GEO:GSM935619 respectively. Mouse ChIP-seq data is from array expression E-MTAB-1414.

## Ethics approval

This study was approved by the Oregon National Primate Research Center (ONPRC) Institutional Animal Care and Use Committee (IACUC) and conforms to current Office of Laboratory Animal Welfares (OLAW) regulations as stipulated in assurance number A3304-01.

## Competing interests

The authors declare no competing financial interests.

## Authors’ contribution

KRC, OV, and DES conceived and designed the experiments. KRC, HS, CT, OV performed experiments. KRC analyzed the data. KRC, HS, OV and DES wrote the paper. The authors have read and approved the final manuscript.

## Supplemental Information

**Supplemental Figure S1.**
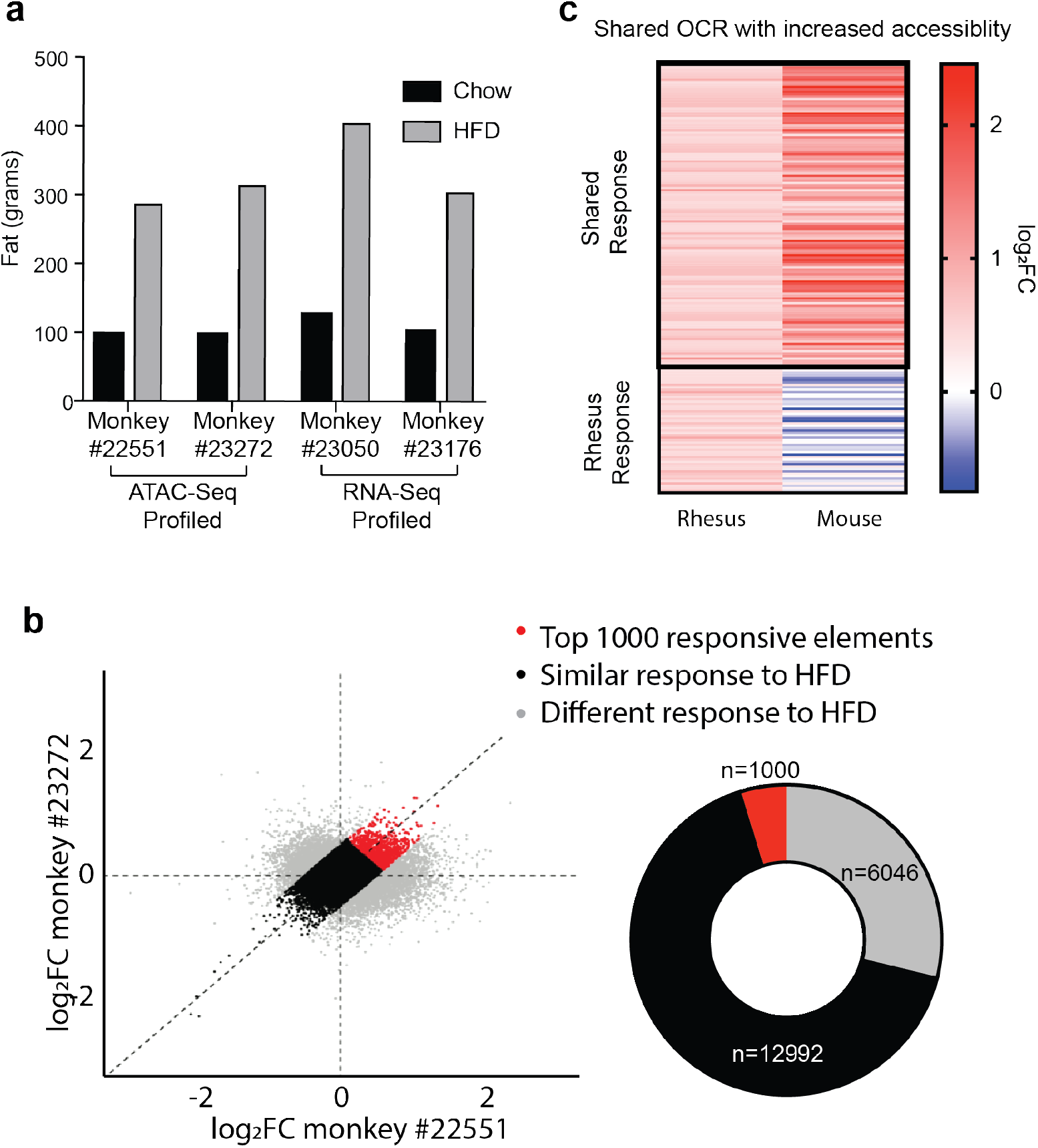
Regulatory element response to high fat diet. **a** Whole body fat (% change relative to pre-HFD) for rhesus macaques fed a standard chow or a high fat diet. **b** selection of regulatory elements with a similar response to high fat diet. Elements with a logFC difference less than 0.50 between the two monkeys were selected to be profiled. The top 1000 responsive elements in both monkeys were selected to be our “responsive” fraction. **c** Comparison of chromatin accessibility changes in response to high fat diet at shared regulatory elements.

**Supplemental Figure S2.**
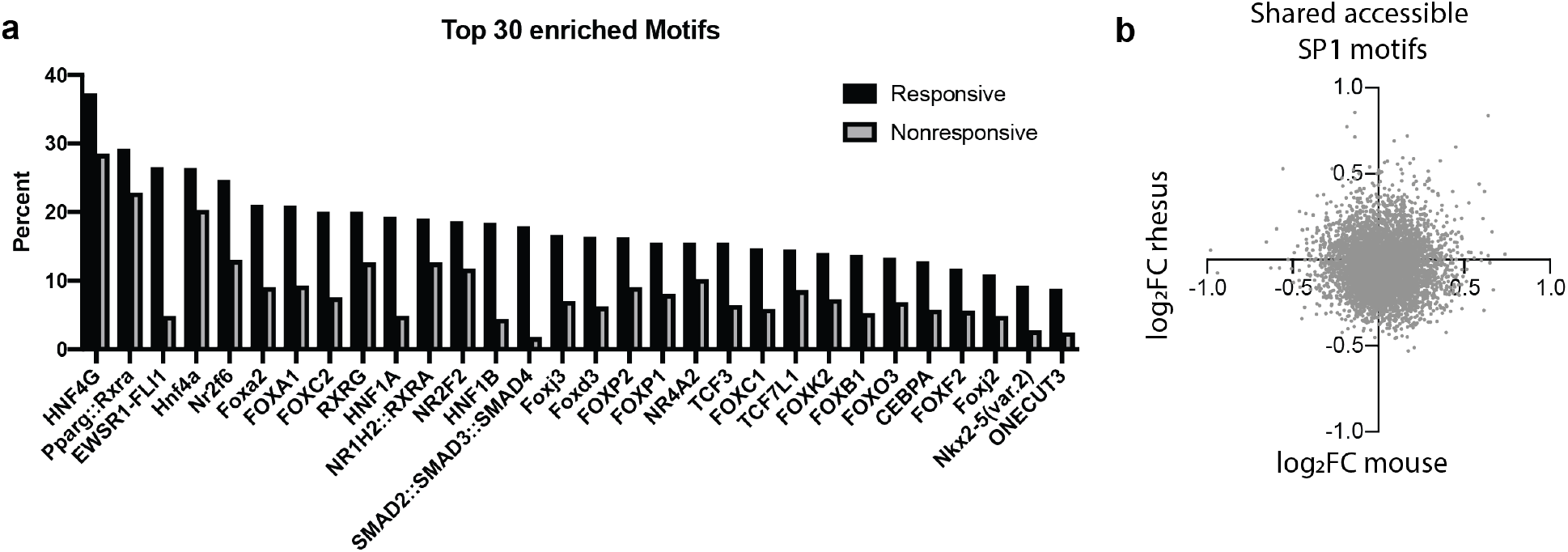
Responsive regulatory element motif profiling. **a** Selection of top 30 motifs enriched in responsive regulatory elements relative to non-responsive elements. **b** Comparison of chromatin response between mice and rhesus macaques at shared regulatory elements with conserved Sp1 motifs.

**Supplemental Figure S3.**
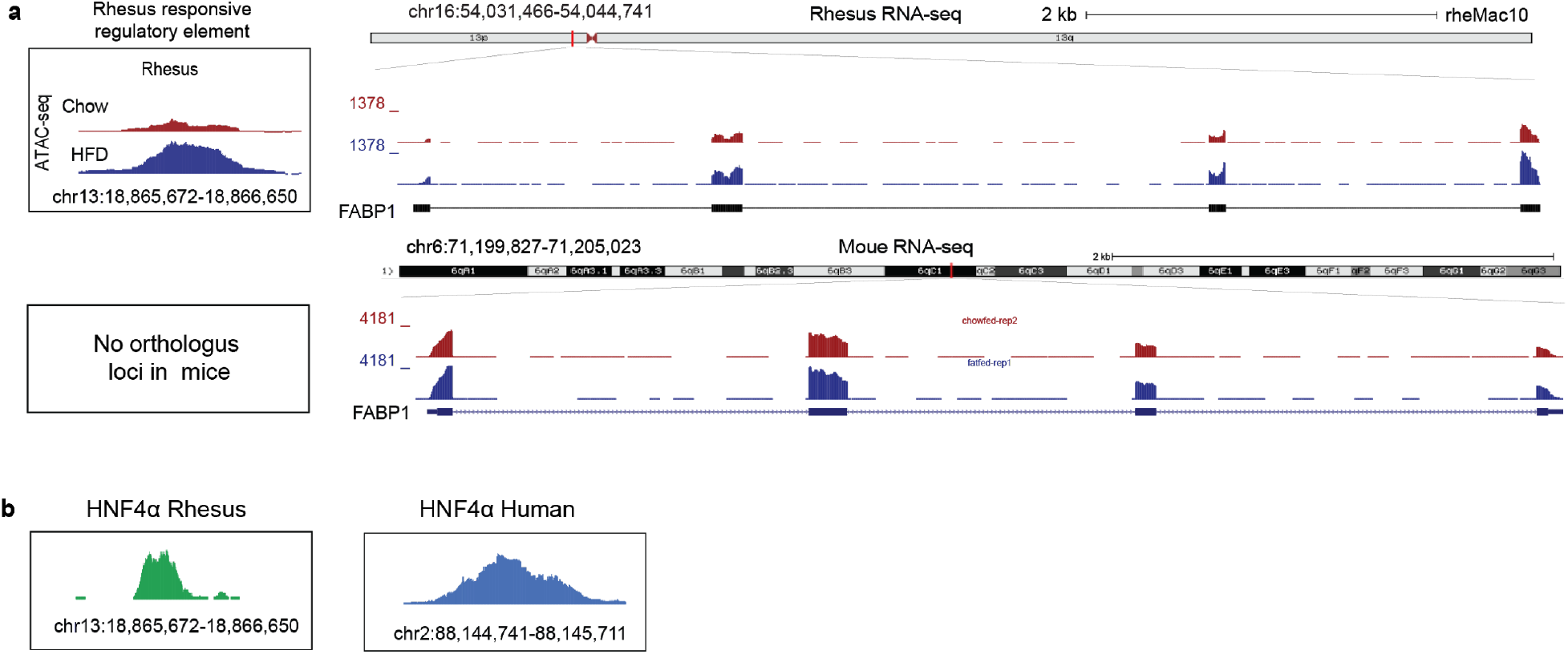
Transcriptional and chromatin response of FABP1 in mice and rhesus macaques. **a** Genome screenshot of RNA-seq (right) and chromatin accessibility (right) of the gene FABP1 and the rhesus responsive regulatory element proximal to its TSSs in both rhesus macaques (top) and mice (bottom). This sequence is only unique to rhesus macaques. **b** ChIP seq signal of HNF4 in both humans and mice at the responsive regulatory element.

*Additional File 2:* Table S1.xls

Physiological changes of rhesus macaques exposed to 6 months of high fat diet.

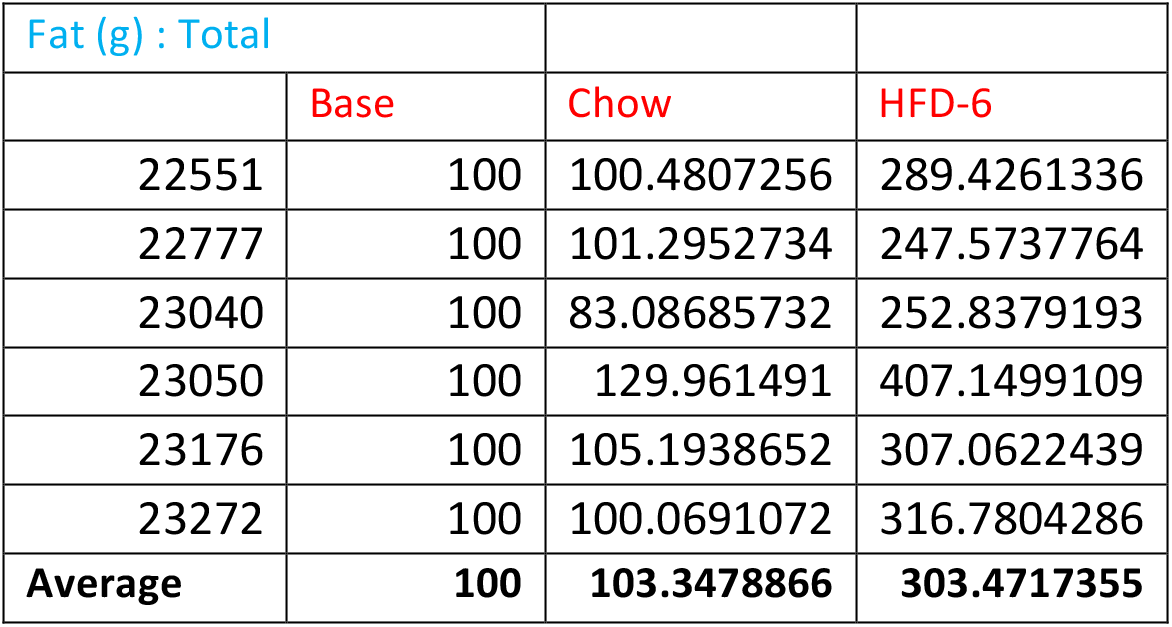
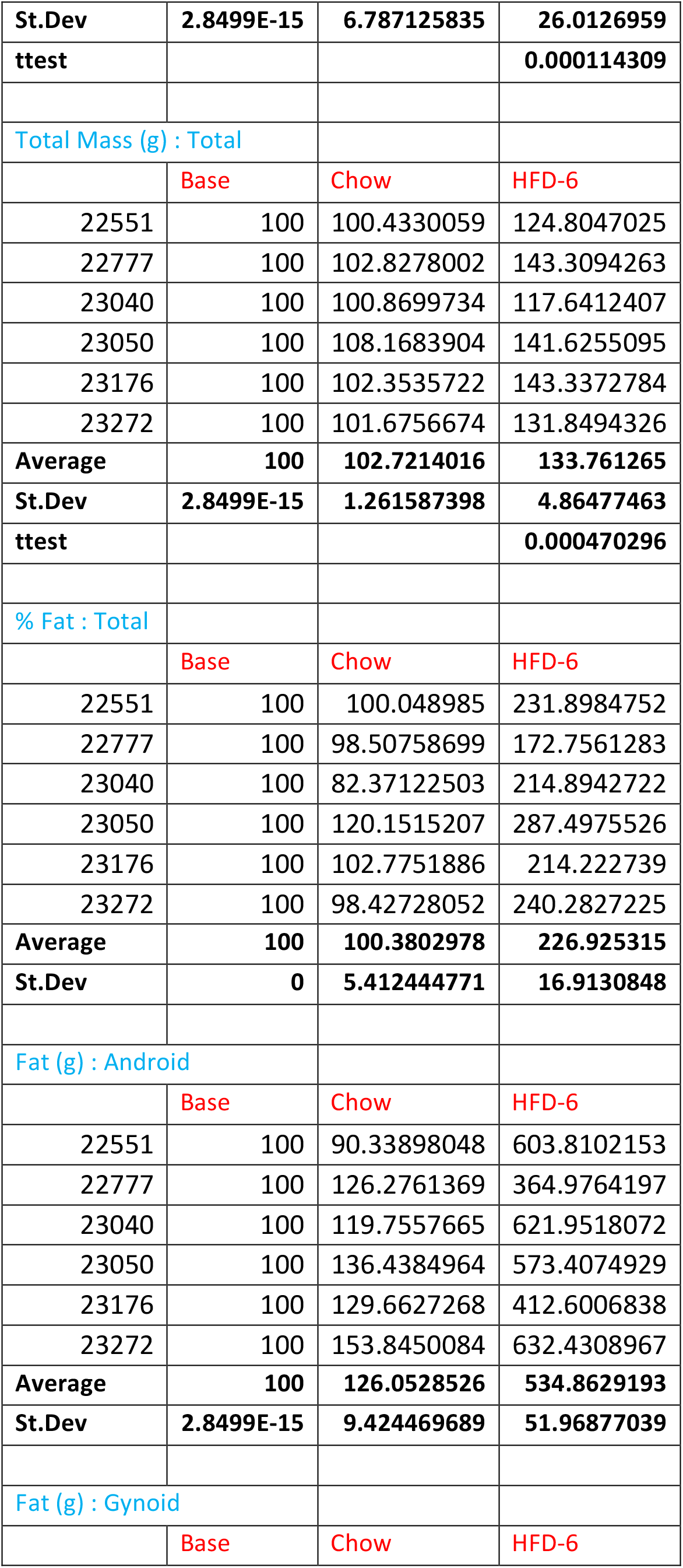
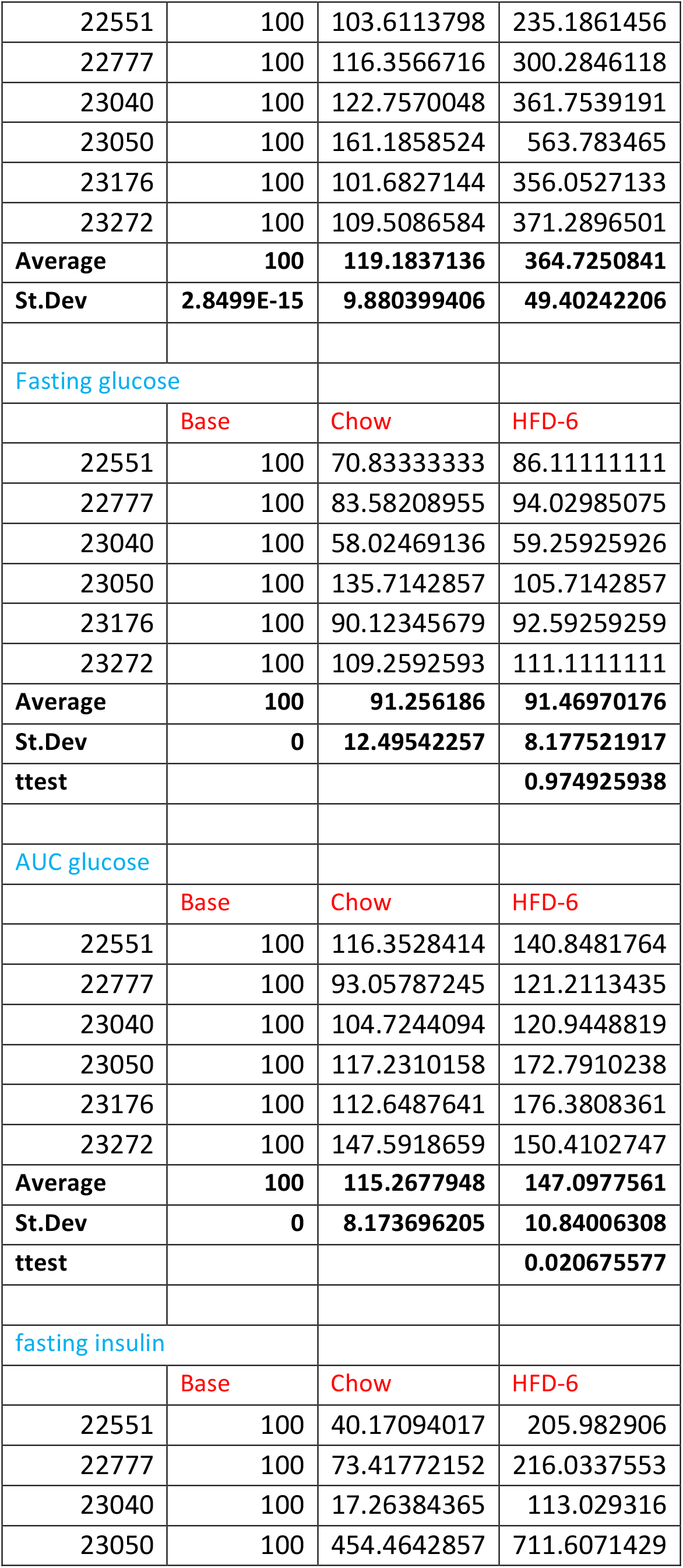
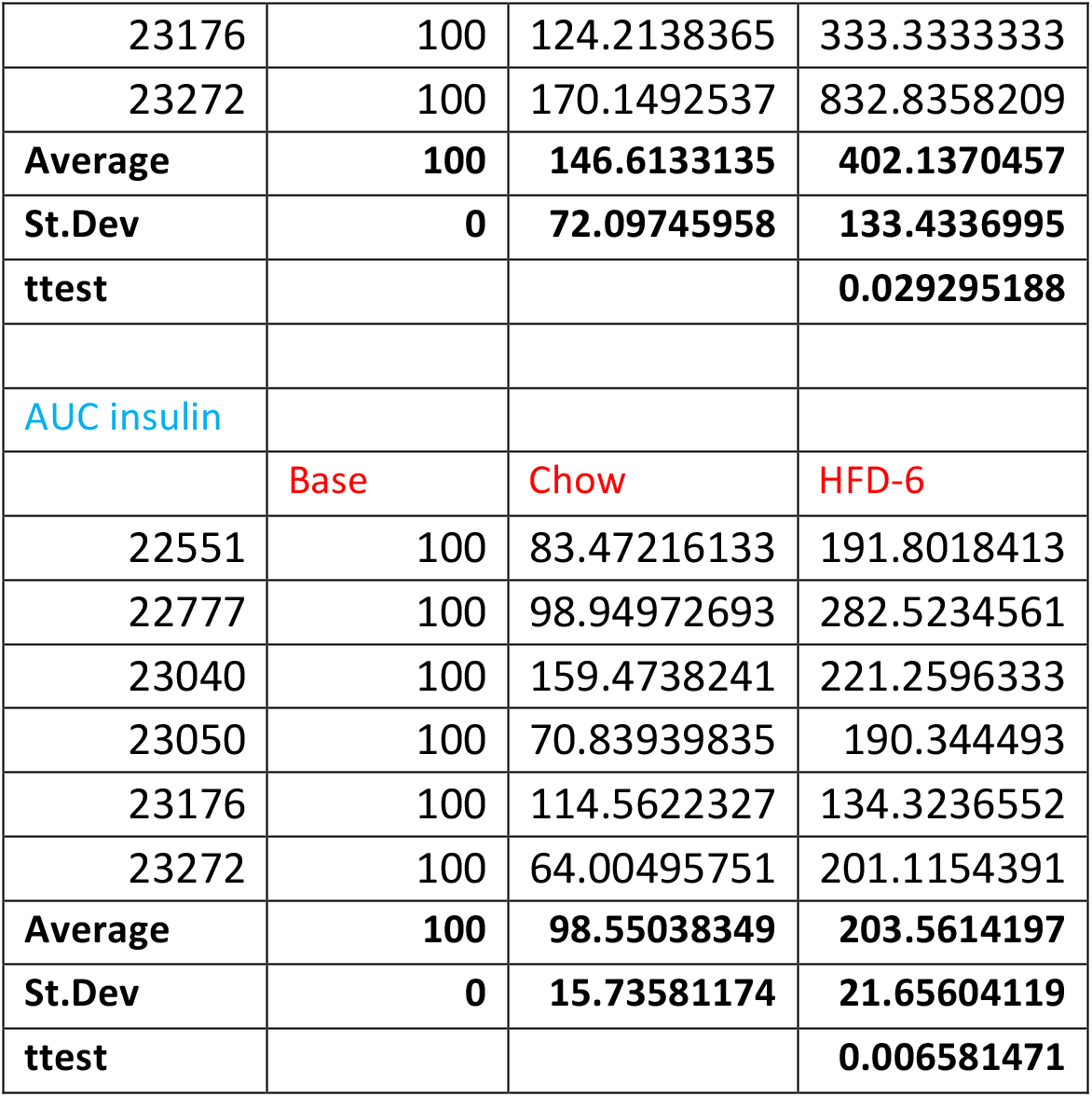

| Fat (g) : Total |  |  |  |
| --- | --- | --- | --- |
|  | Base | Chow | HFD-6 |
| 22551 | 100 | 100.4807256 | 289.4261336 |
| 22777 | 100 | 101.2952734 | 247.5737764 |
| 23040 | 100 | 83.08685732 | 252.8379193 |
| 23050 | 100 | 129.961491 | 407.1499109 |
| 23176 | 100 | 105.1938652 | 307.0622439 |
| 23272 | 100 | 100.0691072 | 316.7804286 |
| Average | 100 | 103.3478866 | 303.4717355 |
| **St.Dev** | **2.8499E-15** | **6.787125835** | **26.0126959** |
| **ttest** |  |  | **0.000114309** |
| Total Mass (g) : Total |  |  |  |
|  | Base | Chow | HFD-6 |
| 22551 | 100 | 100.4330059 | 124.8047025 |
| 22777 | 100 | 102.8278002 | 143.3094263 |
| 23040 | 100 | 100.8699734 | 117.6412407 |
| 23050 | 100 | 108.1683904 | 141.6255095 |
| 23176 | 100 | 102.3535722 | 143.3372784 |
| 23272 | 100 | 101.6756674 | 131.8494326 |
| **Average** | **100** | **102.7214016** | **133.761265** |
| **St.Dev** | **2.8499E-15** | **1.261587398** | **4.86477463** |
| **ttest** |  |  | **0.000470296** |
| % Fat : Total |  |  |  |
|  | Base | Chow | HFD-6 |
| 22551 | 100 | 100.048985 | 231.8984752 |
| 22777 | 100 | 98.50758699 | 172.7561283 |
| 23040 | 100 | 82.37122503 | 214.8942722 |
| 23050 | 100 | 120.1515207 | 287.4975526 |
| 23176 | 100 | 102.7751886 | 214.222739 |
| 23272 | 100 | 98.42728052 | 240.2827225 |
| **Average** | **100** | **100.3802978** | **226.925315** |
| **St.Dev** | **0** | **5.412444771** | **16.9130848** |
| Fat (g) : Android |  |  |  |
|  | Base | Chow | HFD-6 |
| 22551 | 100 | 90.33898048 | 603.8102153 |
| 22777 | 100 | 126.2761369 | 364.9764197 |
| 23040 | 100 | 119.7557665 | 621.9518072 |
| 23050 | 100 | 136.4384964 | 573.4074929 |
| 23176 | 100 | 129.6627268 | 412.6006838 |
| 23272 | 100 | 153.8450084 | 632.4308967 |
| **Average** | **100** | **126.0528526** | **534.8629193** |
| **St.Dev** | **2.8499E-15** | **9.424469689** | **51.96877039** |
| Fat (g) : Gynoid |  |  |  |
|  | Base | Chow | HFD-6 |
| 22551 | 100 | 103.6113798 | 235.1861456 |
| 22777 | 100 | 116.3566716 | 300.2846118 |
| 23040 | 100 | 122.7570048 | 361.7539191 |
| 23050 | 100 | 161.1858524 | 563.783465 |
| 23176 | 100 | 101.6827144 | 356.0527133 |
| 23272 | 100 | 109.5086584 | 371.2896501 |
| **Average** | **100** | **119.1837136** | **364.7250841** |
| **St.Dev** | **2.8499E-15** | **9.880399406** | **49.40242206** |
| Fasting glucose |  |  |  |
|  | Base | Chow | HFD-6 |
| 22551 | 100 | 70.83333333 | 86.11111111 |
| 22777 | 100 | 83.58208955 | 94.02985075 |
| 23040 | 100 | 58.02469136 | 59.25925926 |
| 23050 | 100 | 135.7142857 | 105.7142857 |
| 23176 | 100 | 90.12345679 | 92.59259259 |
| 23272 | 100 | 109.2592593 | 111.1111111 |
| **Average** | **100** | **91.256186** | **91.46970176** |
| **St.Dev** | **0** | **12.49542257** | **8.177521917** |
| **ttest** |  |  | **0.974925938** |
| AUC glucose |  |  |  |
|  | Base | Chow | HFD-6 |
| 22551 | 100 | 116.3528414 | 140.8481764 |
| 22777 | 100 | 93.05787245 | 121.2113435 |
| 23040 | 100 | 104.7244094 | 120.9448819 |
| 23050 | 100 | 117.2310158 | 172.7910238 |
| 23176 | 100 | 112.6487641 | 176.3808361 |
| 23272 | 100 | 147.5918659 | 150.4102747 |
| **Average** | **100** | **115.2677948** | **147.0977561** |
| **St.Dev** | **0** | **8.173696205** | **10.84006308** |
| **ttest** |  |  | **0.020675577** |
| fasting insulin |  |  |  |
|  | Base | Chow | HFD-6 |
| 22551 | 100 | 40.17094017 | 205.982906 |
| 22777 | 100 | 73.41772152 | 216.0337553 |
| 23040 | 100 | 17.26384365 | 113.029316 |
| 23050 | 100 | 454.4642857 | 711.6071429 |
| 23176 | 100 | 124.2138365 | 333.3333333 |
| 23272 | 100 | 170.1492537 | 832.8358209 |
| **Average** | **100** | **146.6133135** | **402.1370457** |
| **St.Dev** | **0** | **72.09745958** | **133.4336995** |
| **ttest** |  |  | **0.029295188** |
| AUC insulin |  |  |  |
|  | Base | Chow | HFD-6 |
| 22551 | 100 | 83.47216133 | 191.8018413 |
| 22777 | 100 | 98.94972693 | 282.5234561 |
| 23040 | 100 | 159.4738241 | 221.2596333 |
| 23050 | 100 | 70.83939835 | 190.344493 |
| 23176 | 100 | 114.5622327 | 134.3236552 |
| 23272 | 100 | 64.00495751 | 201.1154391 |
| **Average** | **100** | **98.55038349** | **203.5614197** |
| **St.Dev** | **0** | **15.73581174** | **21.65604119** |
| **ttest** |  |  | **0.006581471** |

## Notes

### Competing Interest Statement

The authors have declared no competing interest.

